# A temperate *Siphoviridae* bacteriophage isolate from Siberian tiger enhances the virulence of Methicillin-resistant *Staphylococcus* aureus through distinct mechanisms

**DOI:** 10.1101/2021.05.03.442543

**Authors:** Dan Yang, Shuang Wang, Erchao Sun, Yibao Chen, Lin Hua, Rui Zhou, Huanchun Chen, Zhong Peng, Bin Wu

## Abstract

The emergence and worldwide spread of Methicillin-resistant *Staphylococcus aureus* (MRSA) poses a threat to human health. While bacteriophages are recognized as an effective alternative to treat infections caused by drug resistant pathogens, some bacteriophages in particular the temperate bacteriophage may also influence the virulence of the host bacteria in distinct ways. In this study, we isolated a bacteriophage vB_Saus_PHB21 from an epidermal sample of Siberian tiger (*Panthera tigris altaica*) using a MRSA strain SA14 as the indicator. Our following laboratory tests and whole genome sequencing analyses revealed that vB_Saus_PHB21 was a temperate bacteriophage belonging to the *Siphoviridae* family, and this bacteriophage did not contain any virulence genes. However, the integration of PHB21 genome into the host MRSA increased the bacterial capacities of cell adhesion, cell invasion, anti-phagocytosis and biofilm formation. Challenge of the lysogenic strain (SA14^+^) caused severer mortalities in both *Galleria mellonella* and mouse models. Mice challenged with SA14^+^ showed more serious organ lesions and produced higher inflammatory cytokines (IL-8, IFN-γ and TNF-α) compared to those challenged with SA14. In mechanism, we found the integration of PHB21 genome caused the upregulated expression of many genes encoding products involved in bacterial biofilm formation, adherence and invasion to host cells, anti-phagocytosis, and virulence. This study may provide novel knowledge of “bacteria-phage-interactions” in MRSA.

**IMPORTANCE:** The interaction between bacteriophage and bacteria is like a “double-edged sword”: phages can either kill bacteria, or they may contribute to the bacterial fitness and virulence. In general, phages have positive impacts on bacterial fitness and virulence mainly because they carry antimicrobial resistance genes (ARGs) and/or virulence factors encoding genes (VFGs) and they can spread these harmful genes to the host bacteria. However, we found those phages which do not harbor ARGs and/or VFGs may also enhance the bacterial fitness and virulence. In addition, we also found the integration of phage genomes may lead to the upregulated expression of virulence associated genes in bacteria. Our study may provide new insights to redefine the relationship between phage and bacteria, and the results may also remind a cautious way to set phage-therapy for bacterial infections, before which the safety of a phage intends to be used should be fully evaluated.

## INTRODUCTION

The rapid emergence and dissemination of drug-resistant pathogens have posed a big threat to global public health (1). One drug-resistant pathogen of great concern is methicillin-resistant *Staphylococcus aureus* (MRSA), which is a cause of staph infections that are difficult to treat because of resistance to some antibiotics. Since its first description in 1960s, MRSA has become a leading cause of bacterial infections in both health-care and community settings (2). Compared to the infections due to methicillin-susceptible *S. aureus*, MRSA always causes infections with higher mortality rates and results in increased lengths of hospital stays as well as associated health care costs (3). In recent years, even with the ongoing development of new antibiotics, active surveillance efforts and advances in infection prevention, MRSA remains a prominent pathogen with persistently high mortality (4). In 2017, the World Health Organization (WHO) listed MRSA as a pathogen of high priority that urgently requires new therapeutic options (5). In addition to humans, MRSA strains can also persist and cause infections in many animals, including domestic and wild animals (6)

Known as the natural predators of bacteria, bacteriophages (thereafter referred as phages) have been recognized as a good alternative for treating infections caused by drug resistant pathogens since their discovery in 1915 (7). Indeed, phage therapy has achieved a great success in combating drug resistant bacterial infection in particular those caused by multidrug resistant pathogens in several cases during the past decades (8, 9). However, phage therapy still faces some problems, of particular concern is the safety (10). Many phages in particular the temperate phages have been found to carry antimicrobial resistance genes (ARGs) and/or virulence factors encoding genes (VFGs), and they can mediate the spread of these genes, thereby conferring the bacteria antimicrobial resistance and/or increasing bacterial virulence (11, 12). In this study, we isolated a temperate phage that does not carry either ARGs or VFGs from an epidermal sample of a Siberian tiger (*Panthera tigris altaica*) using a MRSA strain as the indicator. However, we found the integration of this phage genome significantly enhanced the virulence of host MRSA strain. To explore the related mechanism, we performed RNA-Seq analysis and we found the integration of this phage genome cased the upregulated expression many genes related to the biofilm formation and virulence in the lysogenic MRSA strain. Our findings will provide novel knowledge of “bacteria-phage-interactions” in MRSA.

## RESULTS

### Isolation and characterization of a MRSA specific temperate phage

Using a MRSA strain SA14 as the indicator, we isolated a bacteriophage designated vB_Saus_PHB21 (thereafter referred as PHB21) from an epidermal sample collected from a Siberian tiger (*Panthera tigris altaica*) in Qingdao Zoo (Qingdao, China) through the conventional double-layer agar method, as described previously (13, 14). Phenotypically, PHB21 formed small round transparent plaques with a clear boundary in the double agar (**Fig. 1A**). Electron microscopy showed PHB21 particles had a rectangular head (length 166 nm ± 3, width 69 nm ± 3) and a long flexible tail (312 nm ± 3) (**Fig. 1B**). These morphological characteristics indicated that PHB21 belonged to the *Siphoviridae* family, according to the latest International Committee on Taxonomy of Viruses (ICTV) classification.

**Fig. 1.**
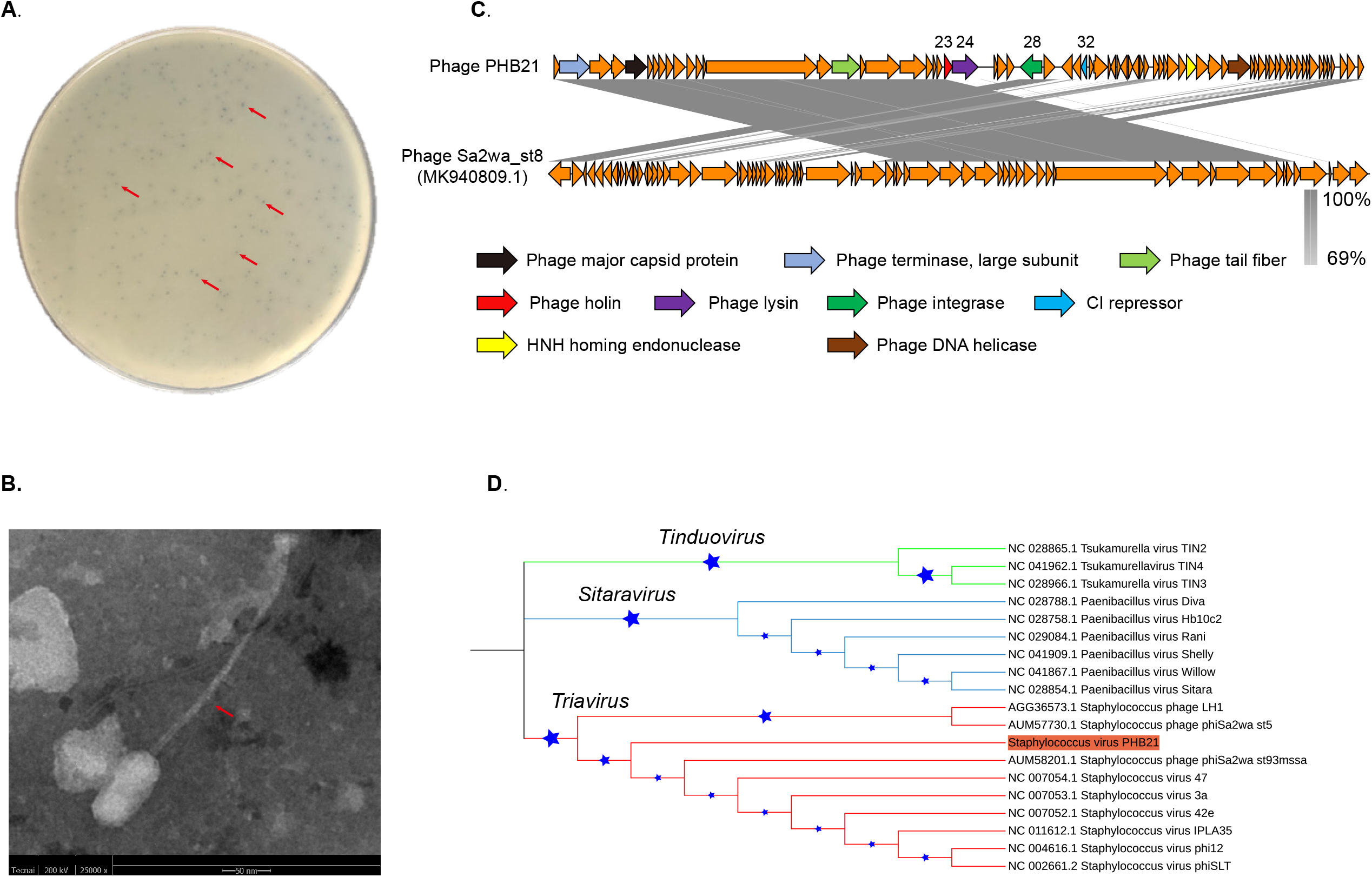
Characteristics of bacteriophage vB_Saus_PHB21. (**A**.) Plaques formed by PHB21 on Methicillin-resistant *Staphylococcus aureus* growing agar; (**B**.) Morphological characteristics of phage PHB21 under the electron microscopy; (**C**.) Comparative analysis of PHB21 complete genome sequence and the complete genome sequence of *Staphylococcus* phage Sa2wa_st8 (GenBank accession no. MK940809.1); arrows refer to the coding sequence regions; (**D**.) Phylogenetic analysis of PHB21 and the other *Staphylococcus* phages; the tree was generated based on nucleotide sequences of the large terminase subunit encoding genes.

Illumina sequencing revealed that PHB21 possessed a linear double-stranded DNA genome with a size of approximately 45.26-Kb in length, with an average G+C content of 33.96%. Sequence alignments revealed that the genome sequence of PHB21 was highly homologous (nucleotide identity 98.57%) to that of a prophage harbored in the genome of *S. aureus* strain NX-T55 (GenBank accession no. CP031839). Bioinformatic analysis showed that PHB21 genome encoded 72 proteins, including a phage integrase (CDS 28) which was highly homologous to that of *Staphylococcus* phage Sa2wa_st8 (GenBank accession no. MK940809.1; **Fig. 1C**), a CI repressor protein (CDS32) which was involved in phage lysogeny process, and an endolysin (CDS24) (**Fig. 1C**). In particular, the genome of PHB21 did not contain any VFGs and/or ARGs. We expressed this endolysin protein in *E. coli* but our laboratory tests showed that the biological activity of this endolysin was not good (date not shown). Phylogenetic analysis based on the nucleotide sequence of the large terminase subunit (ORF1) encoding genes showed that PHB21 was a member of the *Triavirus* species (**Fig. 1D**).

We next performed spot tests to evaluate the lytic capacity of PHB21. A total of 30 MRSA strains of pig or human origin as well as eight strains belonging to the other bacteria species were included for analysis (**Table 1**). The results revealed that PHB21 had a lytic effect on 13 of the 30 MRSA tested; it displayed no lytic effects on *P. multocida, B. bronchiseptica, Salmonella, E. faecalis*, and *E. coli* (**Table 1**).

**Table 1.**
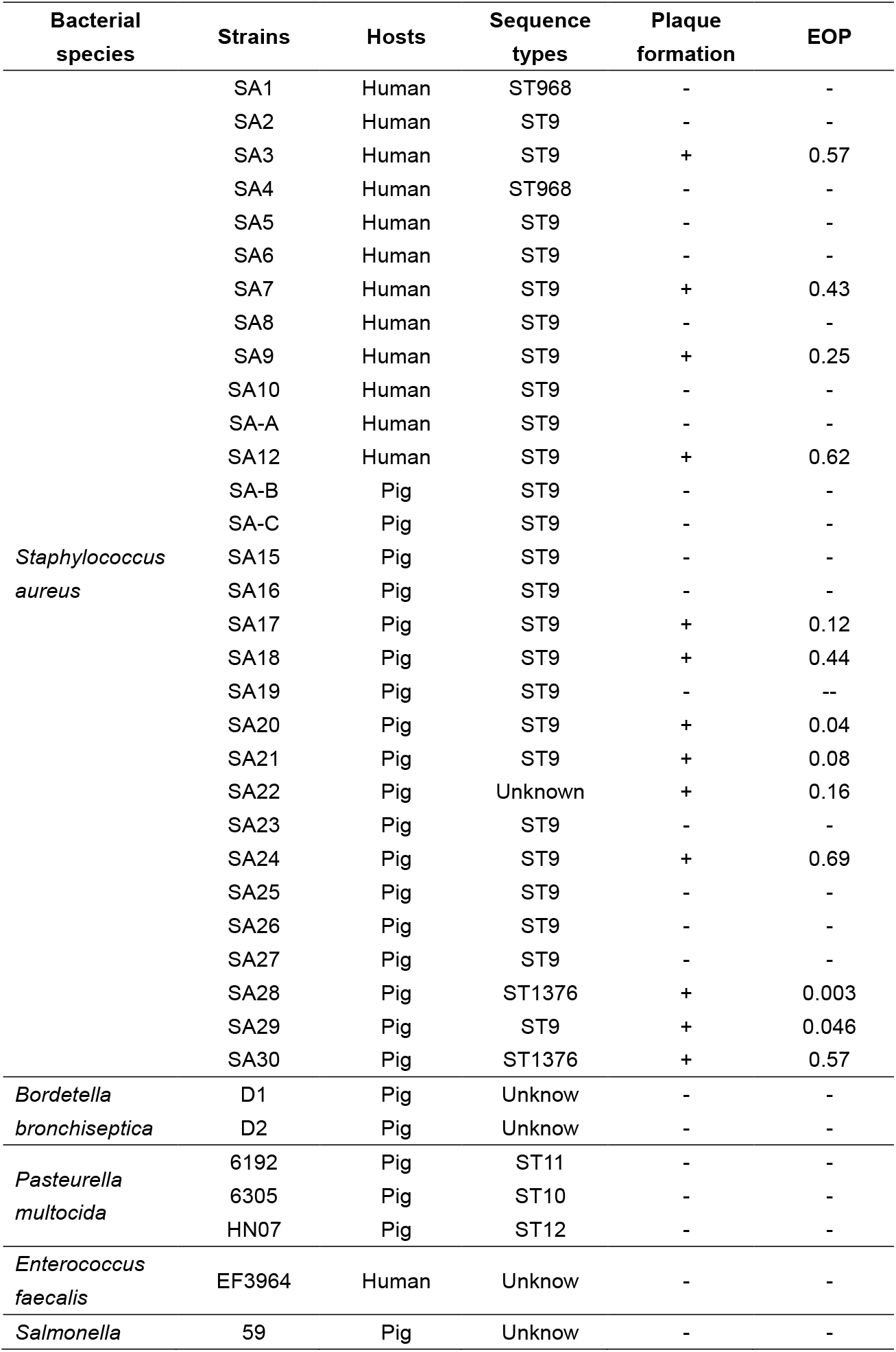
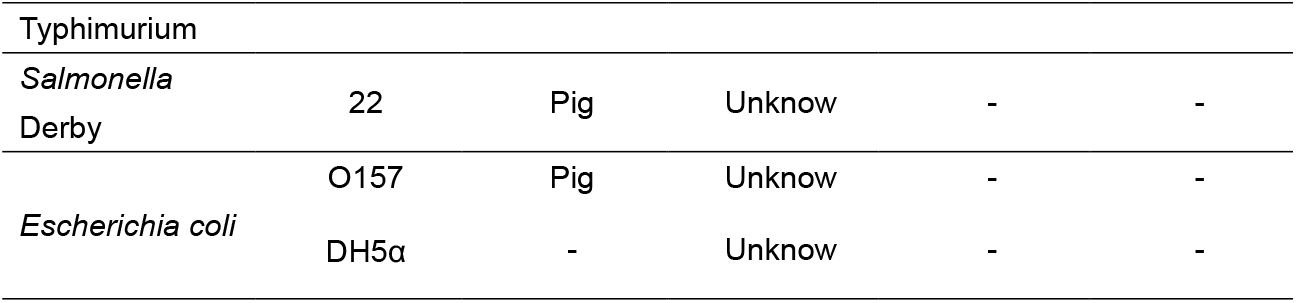
Host spectrum of bacteriophage PHB21

### Determination of phage integration site through ONT sequencing

To determine the integration site of PHB21 genome, we first used this phage to infect the host bacterial strain SA14, and screened the lysogenic strain SA14^+^ (**Figs 2A∼C**). Through Oxford Nanopore Sequencing (ONT), we generated the complete genome sequence of SA14^+^ and the attachment site (attP) of PHB21 genome was determined. We found the attachment site of PHB21 genome was a 29-bp nucleotide sequence “ACCATCACATTATGATGATATGTTTATTT” (**Fig. 2D**). In addition, the phage genome was integrated into a putative coding sequence region (CDS28) of SA14. Annotation through different tools suggested that this coding sequence region encoded a hypothetic protein.

**Fig. 2.**
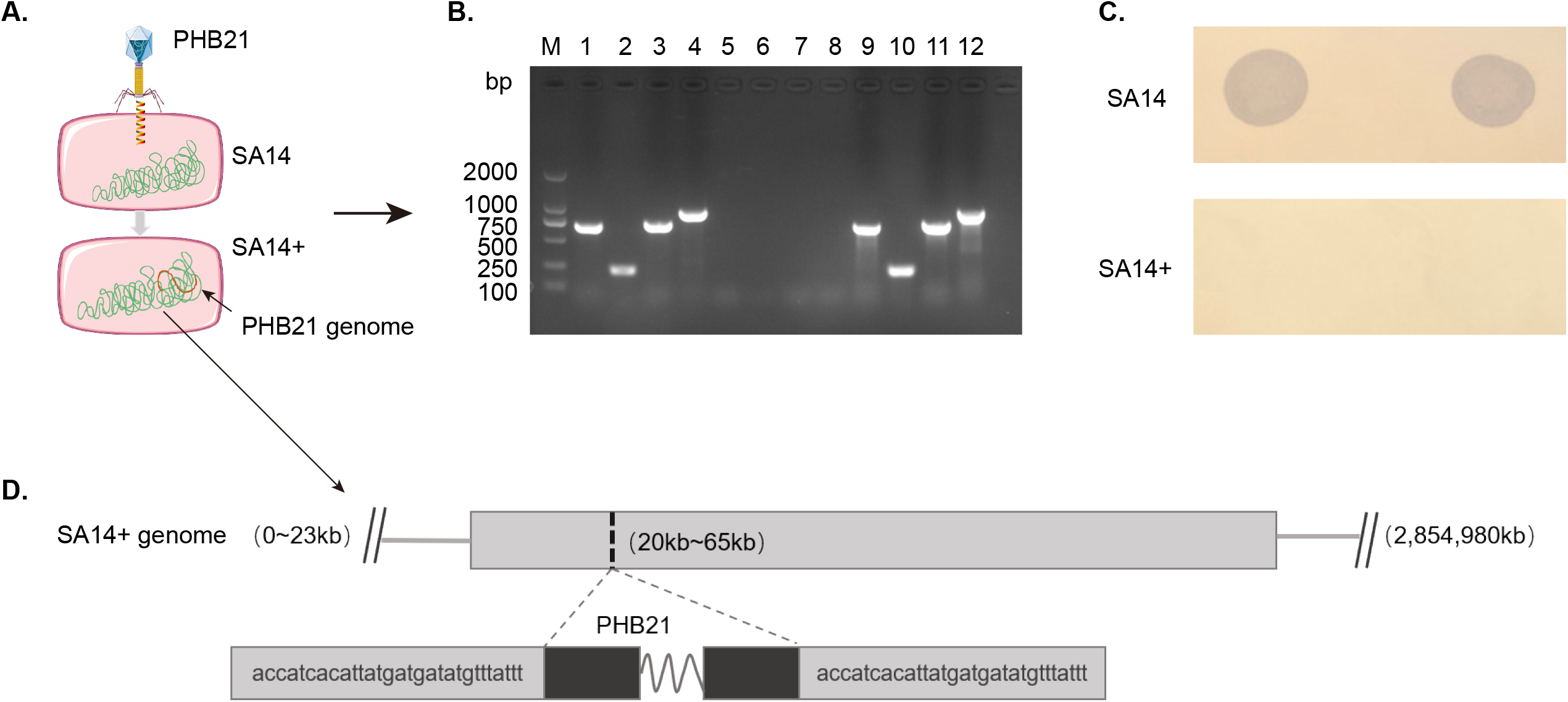
Generation of PHB21 lysogens. (**A**.) A model for the generation of PHB21 lysogens. (**B**.) Lysogenic strains were verified by PCR; M: DL 2000 DNA marker, 1∼4: Using the genomic DNA of MRSA strain SA14^+^ as the template to amplify the phage large terminase gene (band 1; 713 bp), *cI* gene (band 2; 223 bp), major capsid protein gene (band 3; 748 bp) and integrase gene (band 4; 964 bp); 5∼8: Using the genomic DNA of MRSA strain SA14 as the template to amplify the phage large terminase gene (band 5), *cI* gene (band 6), major capsid protein gene (band 7) and integrase gene (band 8); 9∼12: Using the genomic DNA of phage PHB21 as the template to amplify the phage large terminase gene (band 9; 713 bp), *cI* gene (band 10; 223 bp), major capsid protein gene (band 11; 748 bp) and integrase gene (band 12; 964 bp); (**C**.) Phage resistance test; phage PHB21 could lyse MRSA strain SA14, but could not lyse the lysogenic strain SA14^+^; (**D**.) the attachment site (attP) of PHB21 in the chromosome of lyse the lysogenic strain SA14^+^ determined using ONT sequencing.

### Effect of phage-integration on the biological characteristics of MRSA

To explore the influence of the integration of PHB21 genome on the biological characteristics of MRSA strain SA14, we performed a series of laboratory tests. First, we observed the morphologies of both SA14^+^ and SA14, and we did not find any observed differences on bacterial morphology and size between the two strains under transmission electron microscope (**Figs 3A&B**). We then measured the one-step growth curve and tested the bacterial tolerance against mouse and swine sera, and we found SA14^+^ and SA14 showed similar growth curve (**Figs 3C&D**) and capacity of anti-serum bactericidal effects (**Figs 3E&F**). However, during the culture we found the lysogenic strain SA14^+^ displayed a more intense yellow/orange color than those of SA14 (**Fig. 3G**). More importantly, SA14^+^ showed increased biofilm formation and capacities of cell adhesion and invasion, and anti-phagocytosis (**Figs 3H∼M**).

**Fig. 3.**
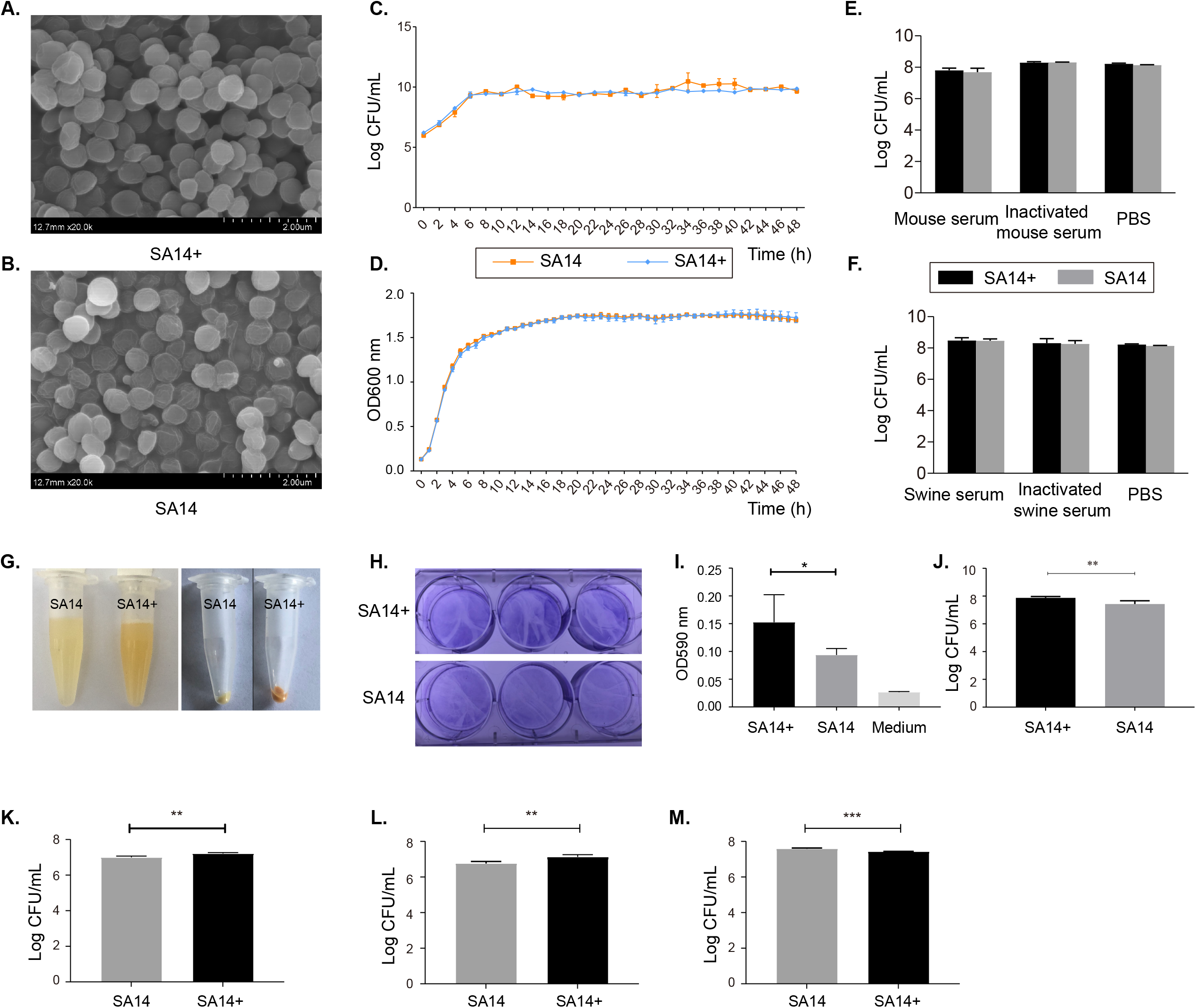
Phenotypical characteristics of the MRSA lysogenic strain SA14^+^ in comparison to those of the MRSA wild type strain SA14. Panels **A** and **B** display the morphological characteristics of SA14^+^ and SA14 under the electron microscopy, respectively; panels **C** and **D** show the growth curves of SA14^+^ and SA14 according to bacterial counting (panel **C**) and measurement of OD_600_ values (panel **D**); panels **E** and **F** exhibit the anti-bactericidal activities of SA14^+^ and SA14 against different types of mouse sera (panel **E**) and pig sera (panel **F**); panel **G** indicates the bacterial color of SA14^+^ and SA14 during the culture; panels **H** to **J** show the biofilm formation of SA14^+^ and SA14 in crystal violet staining (panel **H**), OD_590_ measurement (panel **I**), and bacterial counting (panel **J**); panels **K** to **M** show the capacities of SA14^+^ and SA14 adherence (panel **K**) and invasion to host cells (panel **L**), and anti-phagocytosis (panel **M**). Data represents mean ± SD. The significance level was set at *P* < 0.05 (*).

We next evaluated and compared the virulence of SA14^+^ and SA14 using both *Galleria mellonella* and mouse models. In *Galleria mellonella* models, larvae of *Galleria mellonella* (0.4∼0.5 g per larva, 10 larvae in each of the groups) were challenged with different doses (10^5^ CFU, 10^6^ CFU, 10^7^ CFU, and/or 10^8^ CFU per larva) of SA14^+^ and/or SA14. The number of survival larvae was recoded every 12 h until 144 h. The results revealed that challenge of SA14^+^ induced more severe mortality than SA14 at the same dose (**Figs 4A∼C**). As measured in mouse models, the minimum lethal dose (MLD) of SA14^+^ to mice (BALB/c, 5-week-old) was 1.0 x 10^7^ CFU, which was approximately 7.5-fold lower than that of SA14 (7.5×10^7^ CFU) (**Table 2**). Histologically, mice challenged with SA14^+^ showed more severe lesions in lung, liver and spleen compared to those challenged with SA14 (**Fig 4D**). In particular, challenge of SA14^+^ induced higher levels of IL-8, IFN-γ, and TNF-α in mice (**Figs 4E∼G**).

**Table 2.** Minimum lethal dose (MLD) of *S. aureus* SA14 and SA14^+^ in mouse models

| Strain | Concentration (CFU) | Volume (μL) | Death/Total | Death rate | MLD (CFU) |
| --- | --- | --- | --- | --- | --- |
| SA14 | 1.0×10<sup>9</sup> | 200 | 6/6 | 100% | 7.5×10<sup>7</sup> |
|  | 0.5×10<sup>9</sup> | 200 | 6/6 | 100% |  |
|  | 1.0×10<sup>8</sup> | 200 | 6/6 | 100% |  |
|  | 7.5×10<sup>7</sup> | 200 | 2/6 | 33.3% |  |
|  | 5.0×10<sup>7</sup> | 200 | 0/6 | 0% |  |
|  | 1.0×10<sup>7</sup> | 200 | 0/6 | 0% |  |
|  | 5.0×10<sup>6</sup> | 200 | 0/6 | 0% |  |
| SA14<sup>+</sup> | 1.0×10<sup>9</sup> | 200 | 6/6 | 100% | 1.0×10<sup>7</sup> |
|  | 0.5×10<sup>9</sup> | 200 | 6/6 | 100% |  |
|  | 1.0×10<sup>8</sup> | 200 | 6/6 | 100% |  |
|  | 0.5×10<sup>8</sup> | 200 | 6/6 | 100% |  |
|  | 1.0×10<sup>7</sup> | 200 | 4/6 | 66.6% |  |
|  | 5.0×10<sup>6</sup> | 200 | 0/6 | 0% |  |
|  | 1.0×10<sup>6</sup> | 200 | 0/6 | 0% |  |
| PBS | -- | 200 | 0/6 | 0% |  |
| Negative | -- | 200 | 0/6 | 0% |  |

**Fig. 4.**
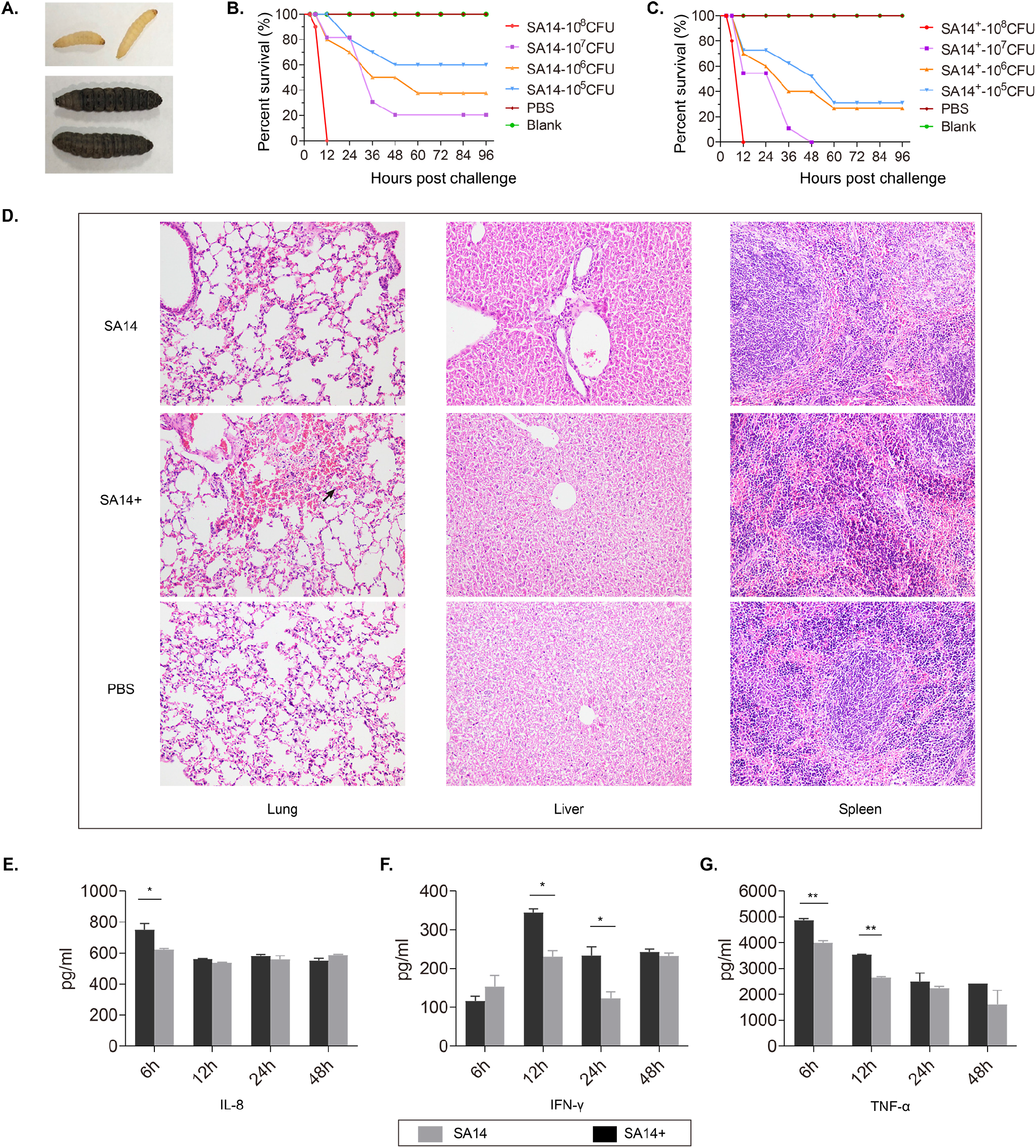
Evaluation of bacterial virulence of MRSA strains SA14^+^ and SA14 using different models. Panel **A** shows the color change of *Galleria mellonella* larvae challenged by MRSA strains; when a larva died due to bacterial infection its color turned to black; panel **B** and **C** display the mortality curves of *Galleria mellonella* larvae due to the infections caused by different concentrations of MRSA strain SA14 and SA14^+^, respectively; panel **D** exhibits the histological damages on mouse lungs, livers and spleens due to the infections of different MRSA strains; panels **E** to **G** show the production of different cytokines (IL-8, IFN-γ, and TNF-α) in mouse sera induced by the infections of different MRSA strains at different time points post challenge. Data represents mean ± SD. The significance level was set at *P* < 0.05 (*).

### Revealing the mechanism of phage enhancing the virulence of MRSA through RNA-Seq

To answer the question why the integration of the non-VFG-carrying phage PHB21 enhances the virulence of the host bacterium, total of RNAs extracted from both SA14^+^ and SA14 at the same time point were sent for transcriptome sequencing. This strategy identified a total of 312 differentially expressed genes (DEGs) in SA14^+^ compared to SA14, including 208 significantly upregulated DEGs and 104 downregulated DEGs (padj < 0.05 & |log2FoldChange| > 0; **Fig. 5A**). In particular, a total of 16 genes associated with the virulence of *S. aureus* exhibited upregulated expressions the lysogenic strain (*clpP1, cap8A, cap8B, cap8C, cap8D, cap8E, cap8F, cap8G, katA, esxA, htpB, clpP2, icaR, cpsK, brkB*, and *clpC*) (**Fig. 5B**).

**Fig. 5.**
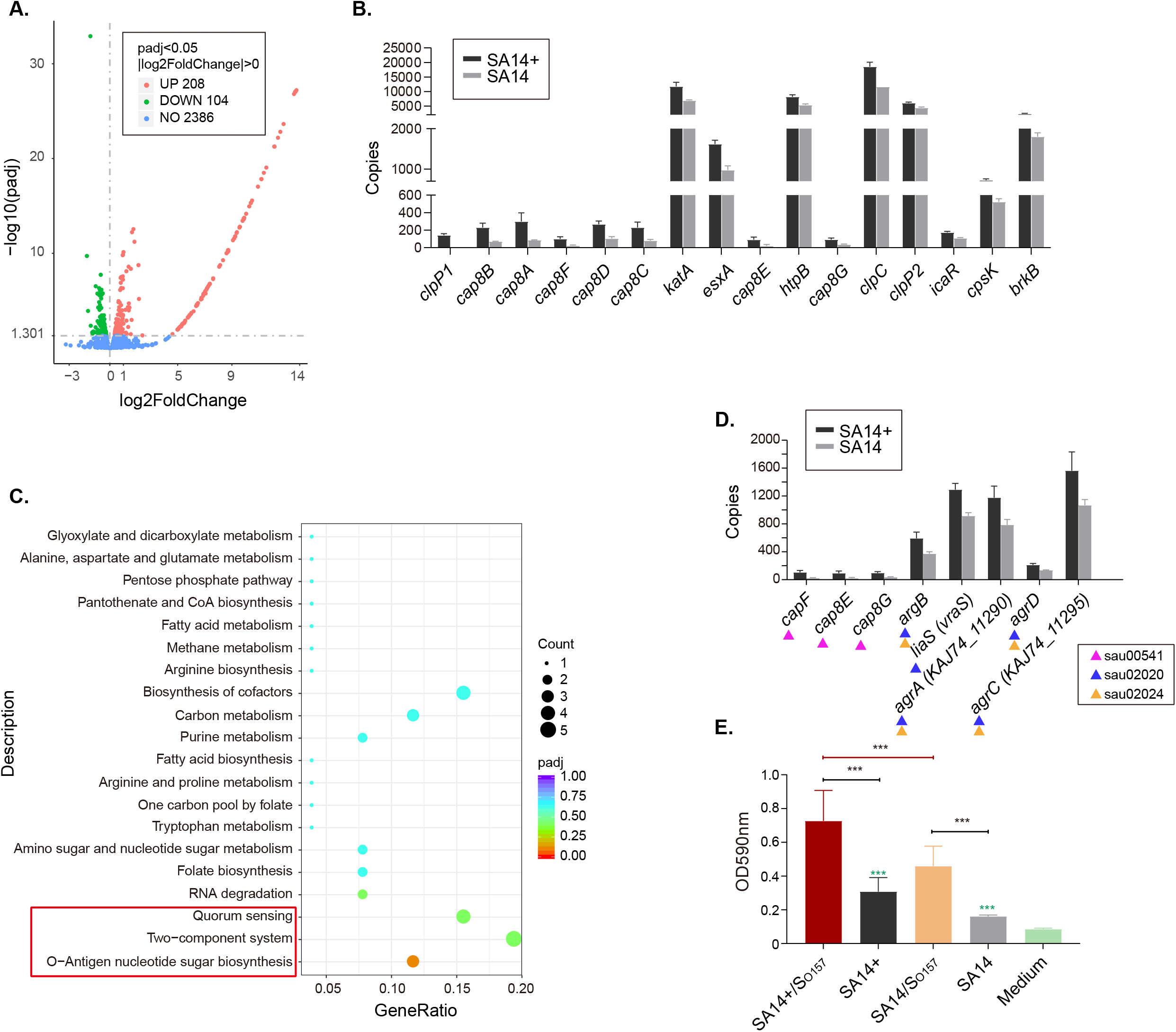
Transcriptome analysis of the MRSA lysogenic strain SA14^+^. (**A**.) Scatter plot displays the differentially expressed genes in the lysogenic strain SA14^+^ compared to those in the wild type strain SA14; (**B**.) Column chart shows the transcription of virulence associated genes in the lysogenic strain SA14^+^ compared to those in the wild type strain SA14; (**C**.) The distribution of KEGG pathways related to the upregulated genes expressed in SA14^+^ compared to the wild type strain SA14; (**D**.) The transcription of genes related to O-Antigen nucleotide sugar biosynthesis (KEGGID: sau00541), two-component system (KEGGID: sau02020), and quorum sensing (KEGGID: sau02024) in MRSA strains SA14^+^ and SA14; (**E**.) Biofilm formation of the lysogenic strain SA14^+^ and the wild type strain SA14 with different treatments. Data represents mean ± SD. The significance level was set at *P* < 0.05 (*).

Kyoto Encyclopedia of Genes and Genomes (KEGG) analysis revealed that the genes upregulated expressed in the lysogenic strain were related to several pathways beneficial for bacterial fitness and pathogenesis, including O-Antigen nucleotide sugar biosynthesis (KEGGID: sau00541), two-component system (KEGGID: sau02020), and quorum sensing (KEGGID: sau02024) (**Figs 5C&D**). Of particular note is the quorum sensing (QS) pathway because it has been reported that bacterial QS signaling has a positive impact on biofilm formation (15, 16). Since bacterial culture supernatant contains QS signal molecules (17, 18), we therefore co-incubated the culture supernatant of *E. coli* O157 and MRSA strains, and measured the biofilm formation. The results revealed that the culture supernatant of *E. coli* O157 enhanced the biofilm formation of both MRSA strains SA14^+^ and SA14 (**Fig. 5E**).

## DISCUSSION

The effect of phages on bacteria is like a “double-edged sword”: phages can either kill bacteria, or they can enhance the bacterial fitness and virulence through different ways (19). Many phages in particular temperate phages have been found to carry VFGs (such as toxin-encoding genes) or ARGs and can spread these harmful genes to bacteria through integration, thereby enhancing bacterial virulence or conferring antimicrobial resistance (11, 12). In this study, the phage PHB21 was isolated from the epidermal sample of a Siberian tiger using a human origin MRSA strain SA14 as the indicator (**Figs 1A&B**). Whole genome sequencing and bioinformatical analysis revealed that PHB21 genome encoded a phage integrase (CDS 28), and a CI repressor protein (CDS32) (**Fig. 1C**). The presence of these two genes suggests that PHB21 is a temperate lysogenic phage since phage integrases mediate unidirectional site-specific recombination between two DNA recognition sequences (20), while CI represses phage lytic genes and commits the phage to the lysogenic program (18). Expectedly, we found the genome of PHB21 could be integrated into the chromosome of the host MRSA strain SA14 through a specific attachment site and the lysogenic strain SA14^+^ was also obtained (**Fig. 2**). However, PHB21 still displayed a capacity to kill several MRSA strains (**Table 1**). This might be because the genomes of these MRSA strains lack the matched attachment sites, and the phage maintain the lytic cycle to those strains (10).

Through ONT sequencing we found PHB21 genome was integrated into a coding sequence encoding a hypothetic protein (**Fig. 2D**). It seems that the disruption of this gene has no effects on the bacterial morphology, growth, and serum-resistance, as we did not see observed differences on these phenotypes of the lysogenic strain SA14^+^ compared to the wild type strain SA14 (**Figs 3A∼F**). However, the lysogenic strain SA14^+^ showed increased yellow/orange in color and biofilm formation (**Figs 3G∼J**). These findings are similar to those from another study (21), and a proposed reason might partly explain the findings is the actions of phage CI protein induce an activation of the alternative sigma factor (SigB) regulon, which provokes an increased secretion of an important virulence factor staphyloxanthin and biofilm formation (21, 22). An increased staphyloxanthin secretion could be indicated by a more intense yellow/orange color of the phage-integration strain during the culture (21). In addition, an increased staphyloxanthin secretion and biofilm formation might also indicate an increased virulence since both of them positively impact the pathogenesis and resistance of MRSA (23). Expectedly, the lysogenic strain SA14^+^ displayed increased cell adhesion and invasion, anti-phagocytosis and virulence to both *Galleria mellonella* and mice compared to those of the host strain SA14 (**Figs 3K∼L, Fig. 4, Table 2**).

Our transcriptional analysis determined a number of genes differentially expressed in the lysogenic strain SA14^+^ compared to those in the wild type strain SA14 (**Fig. 5A**). Of particular note were 16 genes which are important in the pathogenesis of *S. aureus* (**Fig. 5B**). For example, *cap8A, cap8B, cap8C, cap8D, cap8E, cap8F*, and *cap8G* involve in the biosynthesis of capsule, which contributes to the virulence and antiphagocytosis of *S. aureus* (24-26); *clpP1, clpP2* and *clpC* participate in the encoding of Clp chaperones, ATPases and proteases; both of these proteins are central in stress survival, virulence and biofilm formation of *S. aureus* (27, 28); the product of *katA* (catalase enzyme KatA) has a capacity to decompose hydrogen peroxide, which is a reactive oxygen intermediate and is indispensable for the bactericidal activity of phagocytes (29); *esxA* encodes the type VII secretion system (T7SS) effector protein EsxA which is pivotal for bacterial virulence (30); the heat shock protein coding gene *htpB* encodes a 60-kDa chaperonin with an essential function as mediators of protein folding; this protein is important in bacterial pathogenesis and could mediate bacterial attachment to and invasion of host cells (31, 32); *icaR* is a known biofilm regulator gene in *Staphylococcus* and its upregulation contributes to the biofilm formation (33); *cpsK* encodes the teichoic acids export ABC transporter permease subunit TagG which is beneficial for the export of bacterial polysaccharides-wall teichoic acids, thereby contributing to the polysaccharides production (34). In addition, the upregulated-expressed genes induced by the integration of PHB21 genome in the lysogenic strain SA14^+^ are related to several KEGG pathways that have previously been reported to have important roles in the biofilm formation, stress tolerance, and pathogenesis of *Staphylococcus* (21, 22) (**Figs 5C∼E**). The upregulated expression of these genes including the virulence associated genes in the lysogenic strain may explain why the lysogenic strain SA14^+^ displayed increased virulence, biofilm formation, cell adhesion and invasion, and antiphagocytosis compared to the wild type strain SA14 (**Figs 3&4**).

To be concluded, we isolated a temperate MRSA *Siphoviridae* bacteriophage from an epidermal sample of Siberian tiger (*Panthera tigris altaica*) in this study. Although this temperate phage did not carry any virulence genes and still exhibited bactericidal activities to several MRSA strains, it was found to be able to improve the biofilm formation, stress tolerance, and virulence of the lysogenic MRSA strain. This striking phenomenon could be explained through the following reason in mechanism: the integration of the phage genome into the bacterial chromosome led to the upregulated expression of many genes related to the virulence and biofilm formation, which thereby conferring the corresponding phenotypes to the lysogenic strain. Our results presented herein may provide novel knowledge of “bacteria-phage-interactions” in MRSA. This study may also remind a cautious way to set phage-therapy for MRSA infections; the safety of a phage intends to be used should be fully evaluated.

## MATERIALS AND METHODS

### Bacterial strains, phage isolation and purification, electron microscopy, and host specificity

MSRA strains used in this study including those isolates from both humans and pigs in different regions in China (Table 1). These isolates were kindly gifted by Prof. Rui Zhou at Huazhong Agricultural University, Wuhan, China. A human origin MRSA strain SA14 was used as the indicator for bacteriophage isolation. Bacteriophage was isolated and purified according to methods described previously (13, 14). Phage morphology was observed using a 100-kV transmission electron microscope (HITACHI H-7650, Tokyo, Japan) with the same protocol as described by Chen et al. (35). Bacteriophage host range was determined by spot tests according to methods described previously (14). In addition to the MRSA strains mentioned above, our laboratory collected bacterial strains belonging to the other species including *P. multocida, B. bronchiseptica, Salmonella, E. faecalis*, and *E. coli* were also used to determine the host specificity of phage isolated (Table 1).

### Generating PHB21 lysogens

PHB21 (1 × 10^9^ PFU) was spotted on a lawn of MRSA strain SA14 on tryptic soy agar (TSA) plates (Becton, Dickinson and Company, MD, USA) and incubated overnight at 37°C. Bacterial colonies within the lysis zone were isolated and were examined for the sensitivity against PHB21 infection by spot tests. Those with phenotypes of resistance to phage infection were set for PCR amplification of the integrase gene (CDS28), *cI* gene (CDS32), major capsid protein gene (CDS5) and large terminase subunit (CDS2) to confirm the presence of PHB21 genome in the bacterial chromosome.

### Growth test

Overnight culture of SA14^+^ and/or SA14 was transformed into fresh tryptic soy broth (TSB) medium (Becton, Dickinson and Company, MD, USA) at a ratio of 1:1000 (*v*/*v*), which was shaken at 180 rpm, 37 °C. The time point when the shaking culture started was set as 0 h, and from this time point, 100 µL bacterial culture was plated on tryptic soy agar (TSA; Becton, Dickinson and Company, MD, USA) for bacterial counting every 2 h for 48 h. The experiment was repeated three times. In parallel, the same experiments were set to measure the OD_600_ values every 2 h for 48 h by using a fast automatic growth curve analyzer (OyGrowth Curves Ab Ltd, Finland). For both SA14^+^ and SA14, eight repeated samples were measured at each time point.

### Serum bactericidal test, cell adhesion and invasion assay, and anti-phagocytosis test

Bacterial capacity of anti-serum bactericidal activity was evaluated using both mouse and pig sera. Briefly, 25 µL bacterial culture of SA14^+^ and SA14 at mid-log phase were added into 75 µL mouse or pig serum or inactivated mouse or pig serum (inactivated at 56 °C for 30 min). The bacteria-serum mixture was then incubated at 37 °C for 2 h, followed by a series of 10-fold dilutions being performed for bacterial counting. In the control experiment the same volume of PBS replaced the serum. The whole experiment was triplicated.

A previously reported methodology (36) was followed to facilitate the analyses of adhesion and invasion assays, HEp-2 cells (ATCC^®^ CCL-23; approximately 10^6^ cells per well) were infected with MRSA strains SA14^+^ or SA14 at mid-log phase to reach a multiplicity of infection (MOI) of 100:1 (bacteria: cells) and were incubated at 37°C for 2 h. After being washed using PBS for three times, cells were lysed in 1 ml of sterile distilled water. The appropriate diluted lysates were plated on TSA plates to count the adherent and intracellular bacteria. HEp-2 cells were also incubated with ampicillin (100 μg/ml) for 2 hr before lysis. Plating was performed to count the intracellular bacteria alone. All experiments were performed thrice.

Anti-phagocytosis assay was also performed in accordance with previously described methods (36). Briefly, MRSA strains (SA14^+^ or SA14) at log-phase were used to infect RAW264.7 cells to reach a MOI of 10:1. Extracellular bacteria were killed by addition of penicillin (100 μg/ml) with an incubation at 37°C for 2 h. Afterwards, cells were lysed in sterile distilled water and appropriate diluted lysates were plated on TSA plates to count the intracellular bacteria. The experiments were repeated thrice.

### Biofilm formation

Biofilm assays were performed as described previously (21). Overnight cultures of MRSA strains were diluted and 200 µL each of the dilutes poured into the wells of a 96-well microtiter plate (Corning, Corning, NY). The plate was then incubated at 37 °C for 24 h, and the bacterial culture was removed. The wells were washed once using PBS and the floating bacteria were fixed using 200 µL formaldehyde for 30 min at 37 °C. Afterwards, the wells were washed once again using PBS and were stained using 200 µL 1% crystal violet for 15 min. Unbound crystal violet dye in each of the wells was removed and was washed out using PBS. Thereafter, the wells were dried for 2 hr at 37°C, and 100 μl of 95% ethanol was added to each well. Absorbance was recorded at 590 nm using a plate reader (BioTek *Synergy HT*, USA). The wells with a sterile medium were used as negative controls. The experiments were repeated at least thrice.

In another biofilm formation assay, mid-log phase *E. coli* O157 was centrifuged at 8000 rpm for 2 min to discard the pellets. The bacterial supernatant was filtered through an 0.22-µm membrane, and was divided into two samples. One was incubated with 0.2 mg/mL protease K while another one was incubated with PBS both at 56 °C for 30 min. Thereafter, 100 µL of the above *E. coli* supernatant was added into the wells of a 96-well microtiter plate in which contained 100 µL MRSA strains as treated above. The plate was incubated at 37 °C for 24 h, 48 h, 72 h, and 96 h. After incubation, the bacterial culture in different wells was removed and the wells were washed once using PBS. The floating bacteria were fixed using 200 µL formaldehyde at 37 °C, and the wells were washed once again using PBS. Next, the wells were stained using 200 µL 1% crystal violet, and the unbound crystal violet dye in each of the wells was washed out using PBS. Finally, the wells were dried for 2 hr at 37°C, and 100 μl of 95% ethanol was added to each well. Absorbance was recorded at 590 nm using a plate reader (BioTek *Synergy HT*, USA). The wells with a sterile medium were used as negative controls. The experiments were repeated at least thrice.

### Animal tests

Animal tests were performed at Laboratory Animal Center, Huazhong Agricultural University (Wuhan, China), and were approved by the University Research Ethics Committee. All experimental animals were housed under the same atmosphere and were carried out under the guidelines established by the China Regulations for the Administration of Affairs Concerning Experimental Animals (1988) and Regulations for the Administration of Affairs Concerning Experimental Animals in Hubei province (2005).

In *Galleria mellonella* models, each of the 0.4∼0.5 g *Galleria mellonella* larvae received intraperitoneal challenges with MRSA strains at 10^5^ CFU, 10^6^ CFU, 10^7^ CFU, and/or 10^8^ CFU. Control groups included larvae with challenges of PBS and those without any treatment. The number of survival larvae was recoded every 12 h until 144 h. In mouse models, 5-week-old BALB/c mice were divided into 16 groups and each group contained 6 mice. Each mouse in different groups was intraperitoneally challenged with MRSA strains at different doses or PBS or received no treatment (Table 2). After challenge, the mortality was observed and the minimum lethal dose (MLD) was calculated. At 6, 12, 24, 48 hours post challenge mice bloods were harvested and the production of IL-8, IFN-γ, and TNF-α was detected using commercial cytokines detection kits (Dakewe Biotech, Shenzhen, China). Mouse lungs, livers, and spleens were also collected at 7 days post challenge for histological examination.

### Illumina sequencing and Oxford Nanopore sequencing

The genomic DNA of phage PHB21 was extracted using the phenol-chloroform protocol (37). After analysis through electrophoresis on a 1% agarose gel as well as a Qubit 2.0 (Thermo Scientific, Waltham, USA), a NEBNext Ultra™ ‖ DNA Library Prep Kit (NEB, Ipswich, USA) was used to prepare the sequence libraries, which were then sequenced on an Illumina NovaSeq 6000 platform (Novogene Co. LTD, Tianjin, China), using the pair-end 350 bp sequencing protocol. Raw reads with low quality were filtered as previously described (13, 14). High-quality reads were *de novo* assembled using SOAPdenovo2.04 with default parameters (38). Terminal sequences of the phage genome were determined by a previously described modified statistical method (39). Sequence annotation was performed using the RAST Serve (40). Sequence alignments were performed and visualized using Easyfig v.2.0 (41). Phylogenetic trees were generated using MEGAX with 1000 Bootstrap replications (42).

ONT sequencing in combination with the Illumina sequencing was performed to generate the complete genome sequence of the MRSA lysogenic strain SA14^+^. Briefly, the genomic DNA of SA14^+^ was extracted using a commercial bacterial DNA preparing kit (TIANGEN, Beijing, China). DNA quality and quantity were evaluated as described above. Afterwards, an ONT SQK-LSK109 Kit and a NEBNext^®^ Ultra™ DNA Library Prep Kit were used to prepare DNA libraries for ONT sequencing and Illumina sequencing, respectively. Prepared DNA libraries were then sequenced using Nanopore PromethION platform and Illumina NovaSeq PE150, respectively. ONT and Illumina short reads were finally assembled and combined using the Unicycler v0.4.4 software with default parameters. Sequence was also annotated using the RAST Serve (40).

### RNA-Seq

To facilitate transcriptome analysis, MRSA strains SA14^+^ and SA14 were cultured in TSB to mid-log phase (OD_600_ = 0.65). Total RNAs were extracted from each of the samples using Trizol Reagent (Invitrogen, Carlsbad, CA, USA). The quality and quantity of the RNAs isolated were evaluated using NanoPhotometer spectrophotometer from IMPLEN (LA, CA, USA). Sequencing libraries were generated from the qualified RNA samples using a NEBNext^®^ Ultra— RNA Library Prep Kit following manufacturers’ instructions. The libraries were then sequenced on an Illumina Hiseq 2000 platform at Novogene Co. Ltd (Beijing China) and 100-bp paired-end reads were generated. Raw reads obtained from the sequencing were processed using the in-house perl scripts. Next, clean reads with high quality was generated from the raw reads by removing reads containing adapter, reads containing ploy-N, and low-quality reads. Differential expressed genes (DEGs) were determined based on the clean data by using the DESeq R package (1.10.1) with p-value < 0.05 being set. Differential expressed genes were finally annotated by a GOseq R package for Gene Ontology (GO) analysis and a KOBAS software for KEGG analysis. GO terms with corrected p-value less than 0.05 were considered significantly enriched.

### Statistical analysis

Statistical analysis was performed through the “Two-way ANOVA” strategy in GraphPad Prism8.0 (GraphPad Software, San Diego, CA). Data represents mean ± SD. The significance level was set at *P* < 0.05 (*).

## Acknowledgments

We thank staffs at Qingdao Zoo, China for sample collection. This work was supported in part by the Agricultural Science and Technology Innovation Program of Hubei Province (Grant number: 2018skjcx05). Zhong Peng acknowledge the financial support from China Postdoctoral Science Foundation (grant numbers: 2020T130232 and 2018M640719). The funders had no role in study design, data collection and interpretation, or the decision to submit the work for publication.

## Data availability

All sequence data generated in this study are deposited into NCBI database under the Bioproject accession number: PRJNA720778. GenBank accession numbers for the complete genome sequences of *Staphylococcus aureus* bacteriophage phage vB_Saus_PHB21 and *Staphylococcus aureus* lysogenic strain SA14^+^ are MW924497 and CP073012, respectively. Sequence Read Archive (SRA) accession number for the transcriptome sequence is SRR14193019.

